# Climate-driven in-situ trait variation in an annual ruderal grass across Europe

**DOI:** 10.64898/2026.01.19.700245

**Authors:** Helene Villhauer, Timo Hellwig, Sandy Jan Labarosa, Jonathan Moore, Adrian Wysocki, Henryk Straube, Walter Durka, Milan Brankov, Kamil Konowalik, Denise V. M. Blume, Abdenour Kheloufi, Lahouaria Mounia Mansouri, José Manuel Blanco Moreno, Pablo Neira, Joel Torra, Luís P. da Silva, Francesco Santi, Aritz Royo-Esnal, Encarna Rodríguez-García, Carmen Romeralo, Arndt Hampe, Víctor Manzanares-Vázquez, Monika Myśliwy, Anna Bomanowska, Jannis Straube, Nebojša Nikolić, Marta Kolanowska, Ondřej Mudrák, Marcin Nobis, Agnieszka Rewicz, Sabine Metzger, Boris Radak, Sławomir Nowak, Artur Nosalewicz, Anna Elisabeth Backhaus, Katalin Szitár, Miloš Ilić, Severin Einspanier, Izabela Krzemińska, Sylvain Pincebourde, Konrad Kaczmarek, Przemyslaw Baranow, Markus Wagner, Nadine Mitschunas, Albane Bignon, Ali Sahri, Mathieu Leclerc, Agnieszka Nobis, Arne Saatkamp, Andreas D. Drouzas, Anis Edena Zarantonello, Matej Lexa, Stanislav Kopriva, Maria von Korff, Anna Bucharova

## Abstract

1. Plant functional traits link environmental conditions to plant performance and adaptation. Growing evidence shows that in-situ intraspecific trait variation can be as important as differences between species, yet large intraspecific datasets measured in situ are rare. While most studies focused on plant morphological traits, contents of elemental nutrients in the seeds received much less attention so far.
2. We conducted a large-scale in-situ study of the widespread ruderal grass *Hordeum murinum*. We sampled 2070 individuals from 207 populations from a large part of the native range in Europe and Northern Africa. In-situ, we measured seed ripening phenology and morphological traits, and analyzed the content of elemental nutrients in the seeds.
3. We found that *Hordeum murinum* grew larger, produced seeds later, and had heavier seeds in colder and wetter areas. Plants in denser vegetation were taller with heavier seeds but formed fewer spikes. Seed nutrient content generally declined with seed weight and was mainly driven by climate. Soil conditions had only minor effects on plant traits and seed nutrients. Population identity explained much of the variation, indicating a possible genetic component.
4. *Synthesis*: Our findings provide a comprehensive view of *Hordeum murinum* responding to environmental gradients across its European distribution. Climatic variables, especially temperature, are key drivers for reproductive success and seed nutrient content, while local environments, such as biotic pressures, are more critical for growth-related traits. These patterns indicate that *H. murinum* modulates its growth and reproductive investment along environmental gradients, balancing phenology, stress tolerance, and limited competitive capacity.

## Introduction

Plant functional traits allow us to link the environment to plant performance and adaptation (Bruelheide et al., 2018). While most studies have focused on comparing mean functional traits across species or even whole communities (Moles et al., 2009), there is an increasing body of literature documenting intraspecific variation in plant functional traits (De Frenne et al., 2013). This intraspecific variation reflects functional differences between and within plant populations in response to local conditions (Klein-Raufhake et al., 2022), and the magnitude of this variability can be as large as in interspecific comparisons (Tautenhahn et al., 2019). While heritable intraspecific trait variation is typically evaluated in experimental settings (Leiblein-Wild & Tackenberg, 2014), in-situ trait measurements are essential for an ecologically meaningful perspective on how individuals perform in nature (Helsen et al., 2017).

Key life history and morphological traits, such as phenology, plant height, specific leaf area, or seed weight, directly influence plant survival and reproductive success (Adler et al., 2014; Farris & Lechowicz, 1990). Variation in these traits reflects local abiotic and biotic environmental conditions, such as water availability, soil nutrients, pH, or competition (Andrade et al., 2014; Helsen et al., 2017; Lechowicz & Blais, 1988). Life history and morphological trait variation also correlate with large-scale climatic gradients in temperature and precipitation (Leiblein-Wild & Tackenberg, 2014; Lemke et al., 2015). Annual plants growing in warmer and drier climates often flower earlier than their conspecifics growing in colder and wetter climates, which allows them to complete their life cycles before environmental conditions get too dry and hot in summer (Evans et al., 2005). Plants in drier areas generally grow smaller and have lower specific leaf area than conspecifics in wetter regions, a pattern that is interpreted as an adaptation to water scarcity and thermal stress (De Frenne et al., 2013; Poorter et al., 2009). A meta-analysis found that plants from low-latitude populations of the same species produce heavier seeds than high-latitude populations. However, the latitude effects differ considerably among studies (De Frenne et al., 2013). Nevertheless, the data on intraspecific variability in life history and morphological traits are still limited, particularly in high resolution across large geographic and climatic gradients.

Important, but often overlooked, plant traits are the contents of elemental nutrients in the seeds. It is essential for seedling establishment and early survival because young seedlings rely entirely on internal nutrient reserves before root systems develop to acquire nutrients from the soil (Slot et al., 2013). Similar to morphological plant traits, seed nutrient content is also affected by the environment (Sehgal et al., 2018). Nutrient-rich soils generally promote higher seed nutrient contents (Long et al., 2025). Seed nutrient contents also vary along environmental gradients, yet the direction differs between species (Etienne et al., 2018). Most of our understanding of how the environment affects seed nutrients comes from crops, and there is a notable gap regarding the intraspecific variability in the content of elemental nutrients in seeds in natural systems (but see De Frenne, Kolb, et al., 2011; Li et al., 2023; Wang et al., 2025; H. Wu et al., 2024).

Plant traits are not independent of each other, they often covary (Sandel et al., 2016). For example, larger plants typically produce larger seeds (Moles et al., 2004). Relative content of seed elemental nutrients tends to decrease with greater seed weight (Uauy et al., 2006), probably because larger seeds contain a higher proportion of carbohydrates or oils and thus, lower relative content of the other elemental nutrients (Wang et al., 2016). However, empirical data examining intraspecific covariation between seed nutrient content and morphological traits in populations of wild species growing in natural environments remains scarce (De Frenne, Kolb, et al., 2011; Henery & Westoby, 2001)

In-situ intraspecific trait variation in wild plant species is particularly interesting in species with broad geographic distributions. These species encounter a wide range of environmental conditions, which is reflected in a large variation in life history traits (Lemke et al., 2015). Such variation can be related, for example, to climatic, edaphic, or land use factors. However, gradients in these factors often covary across continents, which makes the identification of the drivers of intraspecific variability based on comparison of just a handful of populations difficult (De Frenne et al., 2013; Helsen et al., 2017). Disentangling the effect of individual environmental factors on intraspecific variation thus requires data collected at high resolution on a large scale, not only along the main climatic gradients, but also orthogonal to them. Yet, such data is rare so far.

Here, we focused on a wild grass *Hordeum murinum*, a close relative of cultivated barley. This species is native to Europe and adjacent regions, widespread, and thrives in human-disturbed habitats (Davison, 1977). In-situ, we measured phenology and morphological traits of 2070 plants from 207 populations distributed from North Africa to South Scandinavia. For a subset of 940 plants from 196 populations, we measured the content of elemental nutrients in the seed. We then related these traits to climatic factors, local competition, and soil conditions. We hypothesize that (I) plant phenology, morphological traits, and elemental seed nutrient content covary with each other. (II) Plant phenology, morphological traits, and seed nutrient content vary between in-situ populations across Europe. (III) This variation is affected by local competition, soil properties, and climate.

## Materials and methods

### Study species

*Hordeum murinum* L. is a winter annual grass native to the Mediterranean Region, most of Europe, Central Asia, Western Himalayas, Macaronesia, and the Azores, and has been introduced by humans to Australia, North America, and many other parts of the world, where it became invasive (Fleet & Gill, 2010; Jacobsen & Bothmer, 1995). Throughout most of its range, *H. murinum* typically grows in ruderal areas such as fallow land, roadsides, riverbanks, vineyards, agricultural field margins, and rarely in meadows. It prefers high light intensity, soil nitrogen and phosphorus, and low competition (Davison, 1977).

*H. murinum* flowers typically from early spring to late summer, depending on the region. Flowers are predominantly autogamous, and seeds are dispersed through animals or humans when spikelets adhere to animal fur or human clothing. Seeds typically germinate in late summer or early autumn to winter and overwinter as seedlings or vegetative rosettes (Davison, 1977; Jacobsen & Bothmer, 1995; Mizianty, 2006).

*H. murinum* is a part of an aggregate taxon *Hordeum murinum* agg. that comprises three subspecies differing by cytotypes: diploid (2n = 2x = 14, subspecies *glaucum* (Steud.)), tetraploid (2n = 4x = 28, ssp. *murinum*, or *H. murinum* s. str.), and hexaploid (2n = 6x = 42, ssp. *leporinum* (Link) (Cuadrado et al., 2013).

This study focuses on the tetraploid *H. murinum* subsp. *murinum* in Europe and the Mediterranean (Figure 1). In the southern part of this region, below latitude 45°, *H. murinum* should also occur as diploid (Jacobsen & Bothmer, 1995), but the exact distribution of the two cytotypes is unclear, and we were not able to differentiate the subspecies in the field based on morphology. We thus determined the cytotype of the sampled populations with a flow cytometer (see Appendix). We detected tetraploids throughout the sampled range, and both diploids and tetraploids in Spain (mainland), Algeria, and Morocco. Some populations contained both ploidy levels. As we focus on tetraploids here, we excluded all diploid and mixed populations from further analysis. The in-situ differences between diploids and tetraploids are not the main topic of this study, but they might still be valuable information. Therefore, we included the data alongside certain analyses in the Appendix.

**Figure 1.**
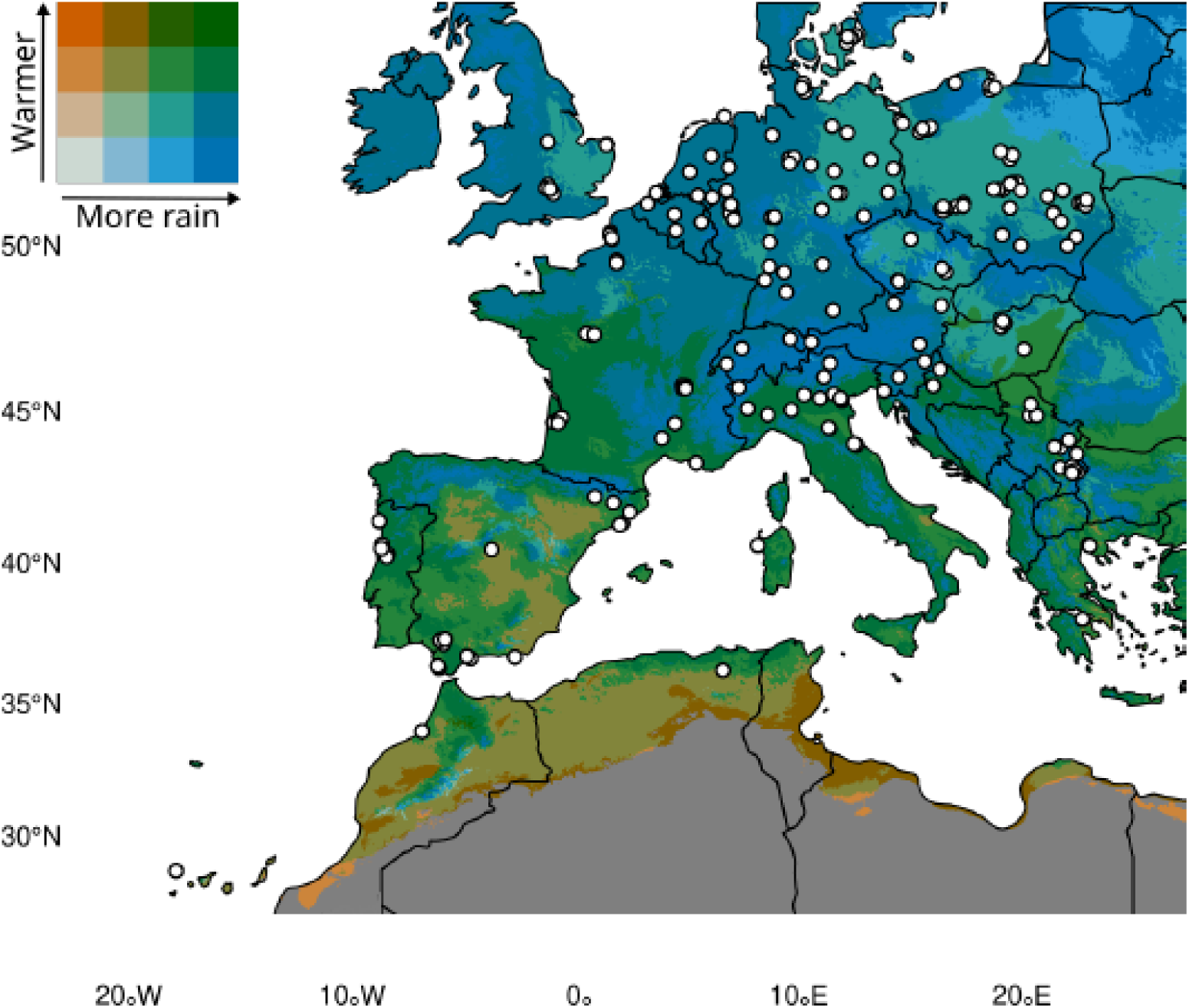
Climate map of the in-situ sampled populations from 2022, 2023, and 2024 of *Hordeum murinum murinum* across Europe and adjacent countries. The map shows the average climate from the years 2000-2020 of mean annual temperature and annual precipitation. It includes only tetraploid populations (excluding diploid and mixed ones).

### Plant material and in-situ trait scoring

We scored and sampled 242 natural populations of *Hordeum murinum L.* across Europe, from which 207 populations turned out to be exclusively tetraploids (Figure 1). Most of the populations were sampled in the summer of 2023, during the seed ripening phase, whereas 10 populations were collected in 2022 (six) and 2024 (four). The seed ripening period varied between May and the end of August, depending on the sampling location. We defined a population as a site with the presence of at least ten individual plants. We kept at least 1 km distance between two adjacent populations to increase the likelihood that the populations are genetically differentiated.

In each population, we randomly collected 10 plants at least 5 meters apart. For each plant, we measured plant height, defined as the longest tiller excluding the spike, and the length of an intact spike (excluding the awns). We then removed the whole plant from the soil, counted the number of tillers with reproductive parts (both unripe and ripe spikes), and harvested one spike that had at least 50 % ripe seeds. In the laboratory, we extracted all seeds from the harvested spikes, counted and weighed them, and stored the clean seeds in a fridge. To standardize the seed weight for further analyses, we extrapolated the weight of a thousand seeds for each plant. For six out of the 207 populations, all spikes were empty with no seeds, but we did obtain morphological and environmental measurements. For two populations, we obtained spikes with ripe seeds and soil samples, but information on plant morphological traits and environmental measurements was missing.

For the following analyses, we excluded the sampling year, as only a few populations were not recorded in 2023, and including the years could distort the data. We also excluded the spike length because spikes lose their seeds as soon as they begin to ripen. Although the sampling protocol required collecting an intact spike, field experience has shown that this was often not possible.

We used the seed collection date as a proxy for the phenology of seed ripening. We mostly sampled the seeds when we could observe the first ripe spikes in a population. Later collection was also possible, because plants typically keep ripe seeds for two to three weeks before complete seed shattering, which introduces some error to this estimate. However, across the whole geographic area, the variation in the seed collection was five months. Thus, the seed collection time captured substantial variability in the onset of seed ripening.

### Environmental variables

To document the environmental conditions of each plant in the field, we estimated vegetation cover and cover of impervious surfaces (e.g., asphalt, cobblestone, concrete) in 1 m² around each sampled plant. We also collected soil samples from beneath each plant and pooled all soil samples for one population. Soil samples were dried at room temperature and stored in the fridge upon arrival at the laboratory.

We obtained climatic information from Chelsa (Karger et al., 2021), which contains data from the years 2000-2020 in 1 km² (30 s) resolution. We downloaded data on nine bioclimatic variables, but because these were highly colinear (r ≥ |0.5|, Figure S1), we included only mean annual temperature (°C), temperature seasonality (°C), and annual precipitation (kg/m²/year) for further analysis.

To measure the pH of the soil, we added a solution of 25 ml of 0.01 M CaCl₂ to 10 g of a soil sample and mixed the two components for one hour. After the sediments had settled, we inserted the electrode of the pH meter (HANNA instruments, pH 211 Microprocessor pH Meter) into the liquid and measured the pH (Rayment & Higginson, 1992).

To obtain plant-available N, P, and S in the soil, we analyzed the soil content of nitrate, phosphate, and sulfate. Specifically, 100 mg soil was extracted in 300 L water for 6 h at 30°C. After 15 min centrifugation at 4°C at maximum speed, the supernatant was transferred into HPLC vials with inlets. Inorganic anions were measured with the Dionex ICS-1100 chromatography system and separated on a Dionex IonPac AS22 RFIC 4x 250 mm analytical column (Thermo Scientific, Darmstadt, Germany; (Dietzen et al., 2020)). 4.5 mM NaCO3/1.4 mM NaHCO3 was used as a running buffer. Standard curves were generated using the external standards of 0.05 mM, 0.1 mM, 0.2 mM, and 0.5 mM KNO3, K2HPO4, K2SO4, and KCl.

### Seed nutrient content

We analyzed the contents of elemental nutrients in the seed (macronutrient Ca, K, Mg, P, S; micronutrients B, Cu, Fe, Mn, Mo, Ni, Zn). We did not analyse content on N, because the selected method does not allow measuring this element (it digests the material in HNO3, see below). We acknowledge that nitrogen is important for plant performance, and the absence of data on this element is a limitation of this study.

To reduce costs, we focused on a subset of six random plants per population and included populations that were at least 10 km apart to capture a broad geographic representation. In total, we analyzed the seed nutrient content of 940 individuals from 196 populations.

The contents of elemental nutrients in the seed were quantified using inductively coupled plasma-mass spectrometry (ICP-MS) (Almario et al., 2017). Two to three seeds (8 to 20 mg) were dried at 60°C overnight and homogenized to a fine powder. The seed material was digested with 500 μL of 67% (w/w) HNO3 in 15 mL Falcon tubes overnight at room temperature. Then, loosely closed tubes containing the samples were placed in a 95°C water bath until the liquid was completely clear (approximately 30 min). After being cooled to room temperature for 10–15 min, the samples were placed on ice, and 4.5 mL of deionized water was carefully added to the tubes. The samples were then centrifuged at 2,000 × g for 30 min at 4°C, and the supernatants were transferred into new tubes. The elemental concentration was determined using an Agilent 7700 ICP-MS (Agilent Technologies). After the analysis, we had to exclude boron (B) from further analysis because the measured values fell in more than 50% of the samples below the detection limit.

### Statistical analysis

First, to understand the strength and direction of covariation among seed ripening onset, morphological traits (height, spike number, seed weight), and contents of elemental nutrients in the seed, we performed a Pearson’s correlation analysis and visualized significant values using a correlation matrix (Figure 2) generated by the *Hmisc* package (Harrell Jr, 2025). Note that seed ripening onset was available at the population-level only, we thus used the population-level average from all traits for the correlation with seed ripening. Correlations among other traits were calculated at the individual plant level. Similarly, we performed a correlation analysis among the environmental predictors (preselected climate variables, vegetation cover, impervious surface, and soil properties). For this, we used population-level averages of each predictor (Figure 3) because most environmental predictors were assessed at the population level, except for vegetation cover and impervious surface. Vegetation cover and impervious surface were strongly correlated with each other (r=-0.62), and this correlation was problematic for the fit of the models below. We thus retained only vegetation cover for further analysis.

**Figure 2.**
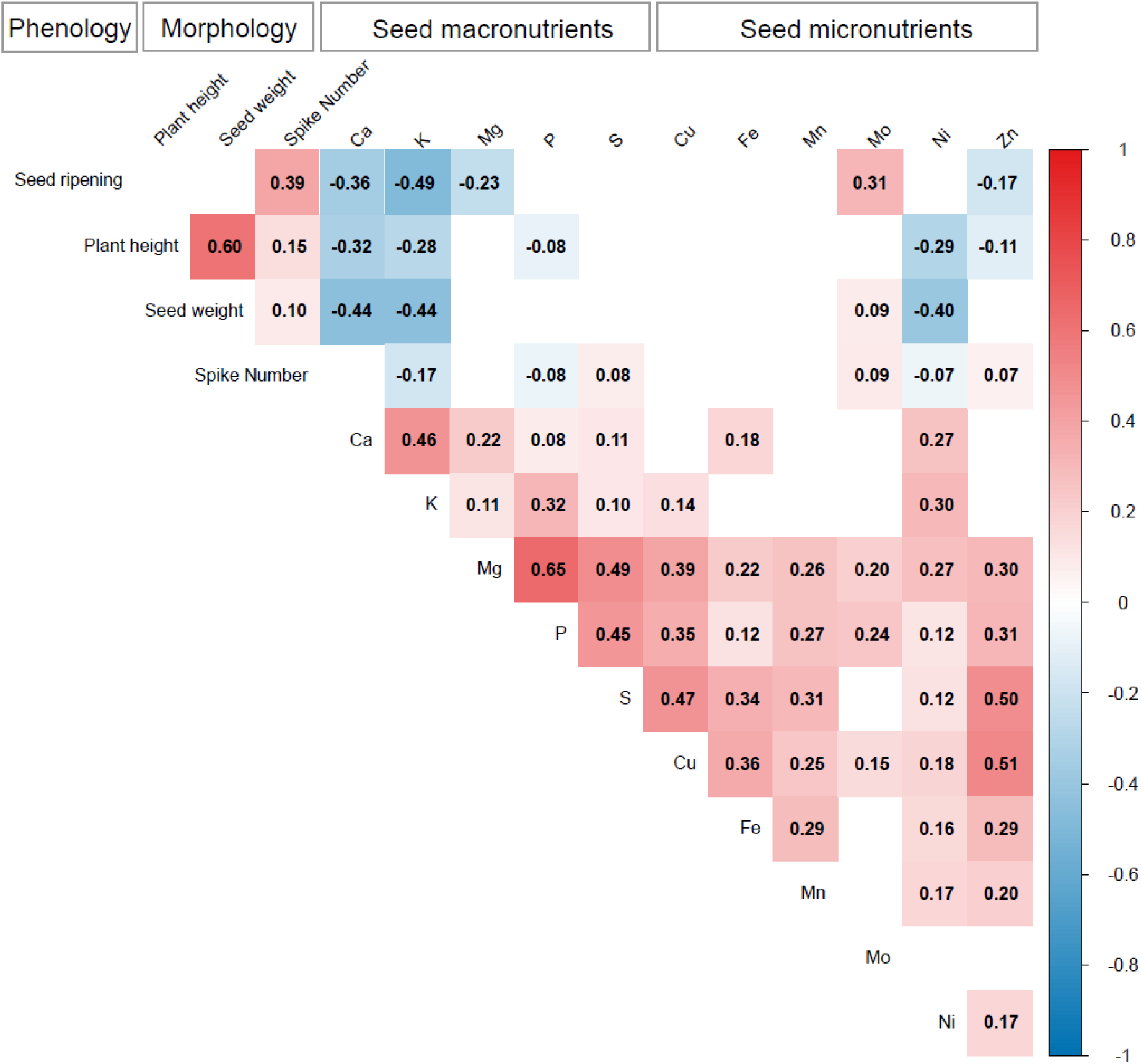
Pearson’s correlation coefficient between plant phenology, morphological traits, and seed nutrient contents (macro- and micronutrients). The number and color intensity in the cells indicate the strength and direction of two related variables. Empty cells indicate non-significant relations (p > 0.05).

**Figure 3.**
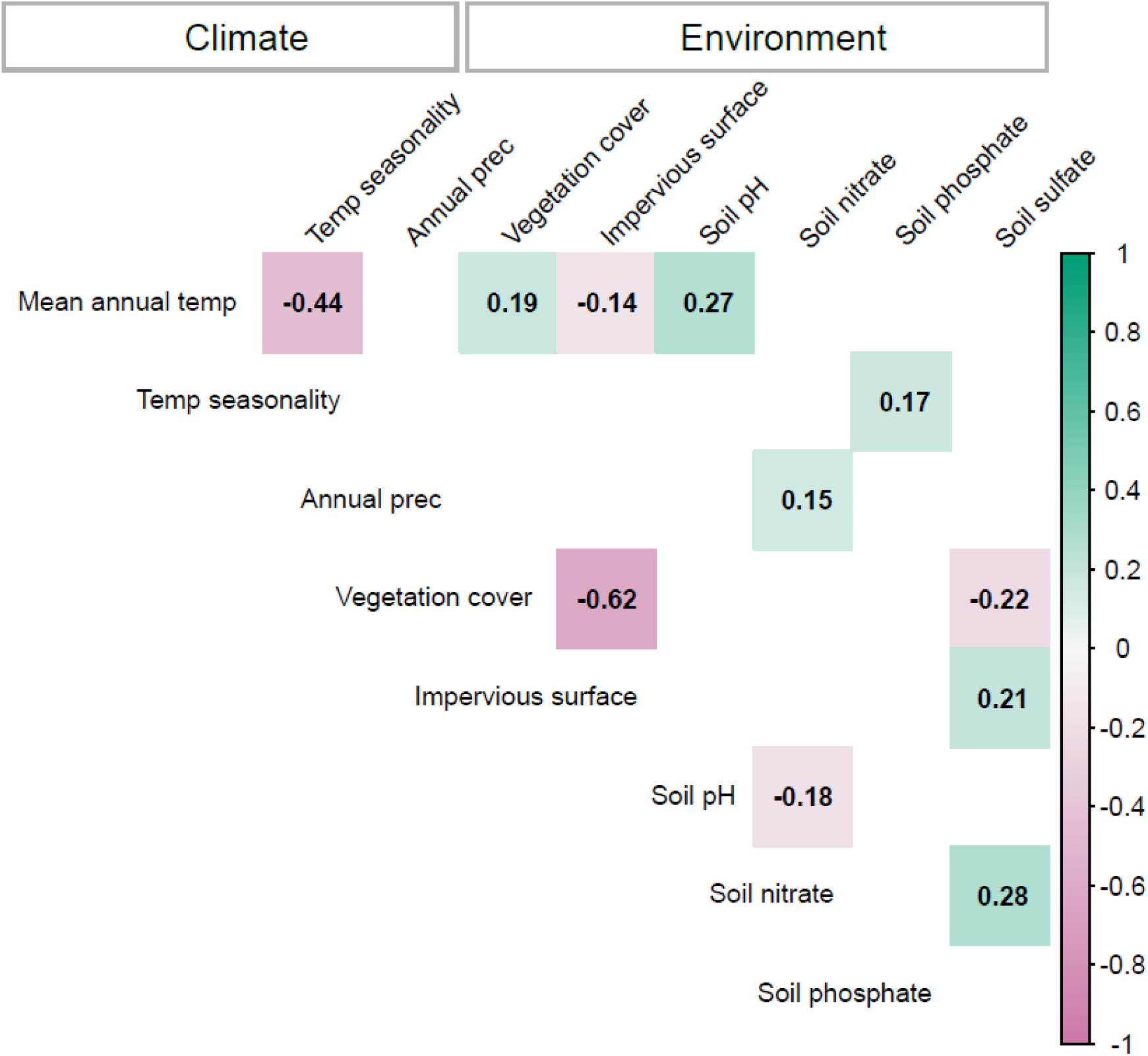
Pearson’s correlation coefficient between all predictors per population. Note that all predictors were averaged across all plants within each population. The number and color intensity in the cells indicate the strength and direction of two related variables. Empty cells indicate non-significant relations (p > 0.05).

Second, we assessed whether morphological traits and the contents of elemental nutrients in the seeds were differentiated among populations. To do this, we related individual plant morphological traits and seed nutrient contents to population identity as a single predictor in a linear model (Table S1). We could not perform this analysis for the onset of seed ripening, because this trait was available on the population level only.

Third, we tested how the onset of seed ripening, morphological traits, and seed nutrient contents related to climate and environmental factors. Explanatory variables were three climatic factors (mean annual temperature, temperature seasonality, and annual precipitation) and five environmental factors (vegetation cover, soil pH, and bioavailable soil nutrients: nitrate, phosphate, and sulfate).

The onset of seed ripening was available only at the population-level. We thus related the onset of seed ripening to population-level averages of the environmental variables in a linear model. To test for spatial autocorrelation in the onset of seed ripening, we applied the function *moran.test*, *spdep* package (Bivand, 2022), on the model and found significant spatial autocorrelation in the ordinary least-squares model residuals (Moran’s I = 0.12, p < 0.001). Therefore, we refitted the model using a spatial lag regression, *spatialreg* package (Bivand et al., 2021), to account for spatial structure (Moran’s I = 0.04, p = 0.08). Because spatial regression models do not provide traditional R² values, we quantified model fit using a pseudo-R² statistic, calculated as one minus the ratio of the residual variance to the total variance of the response variable (Veall & Zimmermann, 1996). To assess the approximate relative importance of each predictor, we compared the changes in the AIC after sequentially removing each predictor from the full model (Burnham & Anderson, 2002). The resulting AIC increases were standardized to percentages to express the relative contribution of each predictor to overall model fit.

Plant morphological traits and contents of elemental nutrients in the seed were available at the level of individual plants. We thus built fourteen linear mixed models, *lme4* package (Bates et al., 2015), one model for each response variable: the three plant morphological traits and the eleven seed nutrient contents. For the fixed explanatory variables, we used the eight climate and environmental variables mentioned above. We included population identity as a random effect to account for the non-independence of plants in one population. To test for spatial autocorrelation, we plotted the residuals from each of the 14 linear mixed models on a map, with point size representing residual magnitude and color indicating whether the residuals were negative or positive. We did not find discernible spatial patterns that would indicate autocorrelation of the residuals and retained the original models (Dale & Fortin, 2014). To determine the proportion of variation explained by the models (fixed, random, and whole model R² value), we used the *rsq* package (Zhang, 2016). To assess the relative importance of individual predictors (fixed effects), we applied the *partR2* package (Stoffel et al., 2021).

To meet model assumptions, we log- or square root-transformed the response variables where necessary (Tables S4, S5, and S6) and evaluated the model fit through residual diagnostics (Zuur et al., 2010). All data analyses were conducted using R version 4.3.1 (R Core Team 2023) within the RStudio environment.

## Results

Reproductive phenology, represented by the onset of seed ripening, was positively correlated with spike number and negatively correlated with the contents of elemental nutrients in the seed (Figure 2). In other words, populations that had ripe seeds later produced more spikes per plant, and the seeds contained lower concentrations of elemental nutrients. Plant morphological traits (plant height, seed weight, and spike number) were all positively correlated with each other, with the strongest correlation between plant height and seed weight (r=0.60), Figure 2. Similarly, seed nutrient contents were in ∼70% of the pairwise correlations positively correlated with each other. The strongest positive correlation was between P and Mg (r=0.65). Seed ripening onset and plant morphological traits were mostly negatively correlated to seed nutrient contents, particularly Ca, K, Mg, P, Ni, and Zn. The strongest negative correlation was between seed ripening onset and K (r=-0.49), and seed weight and Ca and K (both r=-0.44). The only exception was Mo, which was moderately positively correlated with onset of seed ripening (r=0.31), and weakly negatively with seed weight and spike number. Environmental variables (climate, ground cover, soil properties) were correlated with each other only in some cases and only weakly (r<0.3, Figure 3). The only exception was vegetation cover and cover of impervious surface, which were strongly negatively correlated (R=-0.62,), and mean annual temperature and temperature seasonality, which were moderately negatively correlated (r=-0.44, Figure 3).

All three plant morphological traits were significantly differentiated among populations, with population identity explaining 37-38% of the variation (Table S1). The seed nutrient contents were also significantly differentiated between the populations, with the highest differentiation in copper (R^2^=0.57) and the lowest in nickel (R^2^=0.24). Plant morphological traits and seed nutrient contents were also affected by environmental predictors, but these predictors always explained less variability than population identity (pie charts in Figures 4 & 5). Note that we could not assess population differentiation in seed ripening phenology, because we had one value per population.

**Figure 4.**
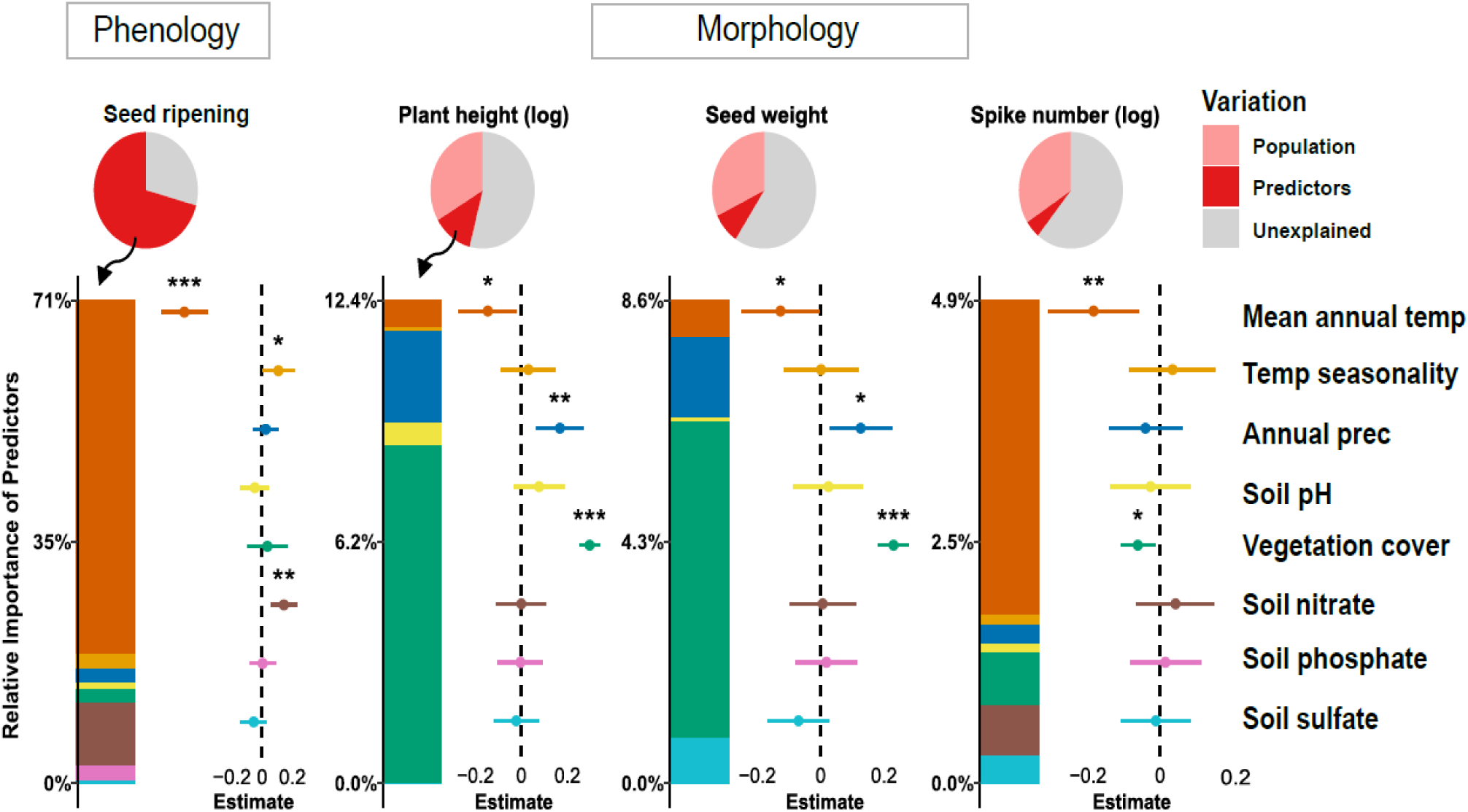
Intraspecific variation in plant phenology and morphological traits. Results of spatial lag model and linear mixed model, respectively, testing for the effect of climate and environment. Each plot contains three main components: (1) *Pie charts* show the R² partitioning of each model split into population (explained variance by the random effect, light red), predictors (explained variance by the fixed effects, dark red), and unexplained variance (grey). (2) *Bar plots* show the estimated relative importance of the environmental predictors (proportion of explained variance of each predictor). (3) *Forest plots* exhibit effect size estimates (points) with confidence intervals (error bars). Asterisks indicate the statistical significance levels of each predictor (* p<0.05, ** p<0.01, *** p<0.001). Predictors were scaled before the analysis. Some response variables were log-transformed before analysis (as indicated in the variable name)

**Figure 5.**
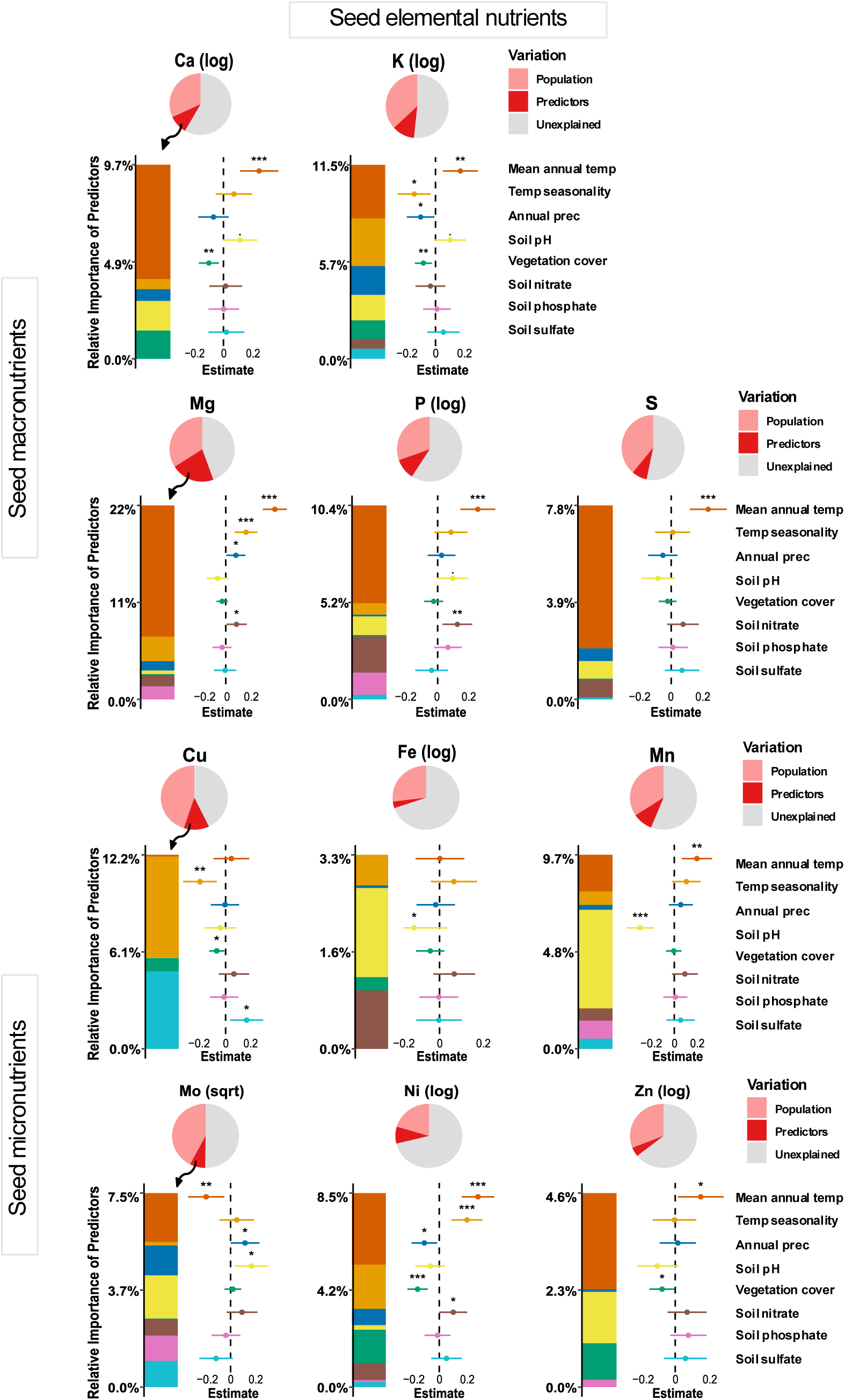
Intraspecific variation in seed elemental contents, results of linear mixed model testing for the effect of climate and environment. Each plot contains three main components: (1) *Pie charts* show the R² partitioning of each model split into population (explained variance by the random effect, light red), predictors (explained variance by the fixed effects, dark red), and unexplained variance (grey). (2) *Bar plots* show the estimated relative importance of the environmental predictors (proportion of explained variance of each predictor). (3) *Forest plots* exhibit effect size estimates (points) with confidence intervals (error bars). Asterisks indicate the statistical significance levels of each predictor (* p<0.05, ** p<0.01, *** p<0.001). Predictors were scaled before the analysis. Some response variables were log-transformed before analysis (as indicated in the variable name).

Seed ripening phenology was primarily affected by climate and nitrate content in the soil. Specifically, plants from areas with higher average temperatures and lower temperature seasonality had ripe seeds earlier. These two climate variables explained about 88% of the variability in seed collection timing among populations. Seeds were also ripe later in soils rich in nitrate (R²=0.04).

Plant morphological traits were affected by vegetation cover around the plant and by climate. Plant height and seed weight were primarily affected by vegetation cover, annual precipitation, and mean annual temperature, with increased vegetation cover, higher annual precipitation, and lower mean annual temperature leading to taller plants and heavier seeds (Figure 4). The number of spikes was most affected by mean annual temperature and vegetation cover, with spike numbers being higher in colder climates and when plants were surrounded by less vegetation.

The contents of elemental nutrients in seeds were most affected by the mean annual temperature (Ca, K, Mg, P, S, Mn, Mo, Ni, and Zn). Specifically, the warmer the climate, the higher the nutrient content (Figure 5), except for Mo, which showed an opposite pattern. The effect of temperature seasonality and mean annual precipitation varied between nutrients, for example, K and Cu decreased with increasing seasonality, Mg and Ni increased. The effect of non-climatic predictors also varied among individual seed nutrients. For example, seed Fe and Mn contents significantly decreased with soil pH, while Mo increased with soil pH. Nitrate had a positive effect on the content of Mg, P, and Ni in seeds. Other soil properties, such as phosphate and sulfate levels, had mostly no effect on the micronutrient content in the seeds, except for soil sulfate, which had a positive effect on seed Cu. Vegetation cover had a negative effect on seed Ca, K, Zn, Cu, and Ni, but the variance explained by this variable was low (Figure 5).

## Discussion

To understand how plants respond to environmental changes and predict range shifts, it is vital to study plant traits in combination with local environmental and climatic factors (Moran et al., 2016). Here, we present a comprehensive, continental-scale dataset on intraspecific variability of a wild species, spanning 4500 km. The dataset includes in-situ seed ripening phenology and morphological measurements of 2070 individual plants from 207 populations, and is complemented by contents of elemental nutrients in the seeds for a subset of 940 individuals of *Hordeum murinum*, an annual ruderal grass species. To the best of our knowledge, this is the first dataset of its kind for a wild plant species under natural conditions, particularly with respect to seed nutrient profiles.

Our results reveal clear ecological patterns: *Hordeum murinum* in colder and wetter areas grew larger, produced seeds later, and had heavier seeds. Plants growing in denser vegetation were also taller and had heavier seeds, but produced fewer spikes. The concentration of some nutrients in seeds declined with seed weight and was mainly affected by climatic factors. Surprisingly, soil condition had only a minor effect on both plant morphological traits and seed nutrient content. Importantly, population identity explained a substantial proportion of trait and seed nutrient variation beyond environmental factors, suggesting that part of the observed variation is due to genetic differentiation between populations.

### Plant phenology and morphological traits

Plants in colder regions had ripe seeds later in the season. Specifically, seed collection date and thus, seed ripening time varied between March in Mediterranean areas and the beginning of September in northern Central Europe. The temperature effects on plant phenology are ubiquitous among plants (Boyko et al., 2023). Particularly in annuals, plants in warm and dry regions commonly complete their life cycle earlier and senesce before summer drought and heat to avoid losses (Evans et al., 2005). Our results are in line with this common phenomenon

Plants were generally larger in colder and wetter regions. Specifically, plant height, seed weight, and number of spikes were all positively intercorrelated and decreased with mean annual temperature, and increased with mean annual precipitation (significant for the first two traits). This contrasts with the pattern observed across species that higher temperatures, typically associated with increased solar radiation, promote biomass accumulation and the production of larger seeds (Murray et al., 2004). However, at the intraspecific level, the relationship between plant height and seed weight and mean annual temperature is highly variable among species, even within grasses (De Frenne et al., 2013; Griffin-Nolan & Sandel, 2023). A similar increase in plant height in colder areas has been reported by De Frenne et al. (2011), who attribute this to the longer photoperiod during the growth season in northern (colder, longer photoperiod during growing season) versus southern (warmer, shorter photoperiod) parts of Europe. In the case of *Hordeum murinum*, plants in the warmer and drier regions in the south senesce in spring, which means they grow during winter when the photoperiod is short. On the other hand, plants in the colder and wetter regions can prolong their growth into late spring when the photoperiod is long, accumulate more biomass and consequently more carbohydrates in the seeds (Dupont & Altenbach, 2003). Plant height and seed weight also increased in denser vegetation (Figure 4). Plants commonly grow taller in denser vegetation to escape competition (Moles et al., 2009), and plant size is positively genetically correlated with seed weight in barley (He et al., 2023). Dense vegetation also weakly reduced the number of spikes, probably because the plant invested more in elongation growth to escape competition than in producing additional flowers. Plants in denser vegetation thus invested their carbohydrates in fewer seeds, which relatively increased the seed weight (Salisbury, 1942). It is also possible that denser vegetation implies more resources that can be invested in the seed, but vegetation cover correlated neither with precipitation as a proxy of water availability, nor with nutrient content in the soil, except for sulphate, but this relationship was weak.

Available soil nutrients and soil pH had no significant effect on the three plant morphological traits, and only a limited effect on plant phenology. This is surprising given the ample literature documenting the effect of nutrient addition on size-related plant traits like height or number of flowers (Andrade et al., 2014; Barker & Pilbeam, 2015; Lechowicz & Blais, 1988). One possible reason may lie in our sampling design: we used one bulk soil sample per population. If the soil conditions of individual plants within the site strongly varied, we did not capture this variation, which may have reduced our ability to detect the effect of soil on plant performance. On the other hand, our results correspond to recent analyses of global intraspecific trait variability in grasses, which have found that soil characteristics are far less important than climate (Griffin-Nolan et al., 2025).

### Content of elemental nutrients in the seeds

Our dataset, comprising seeds from almost 1000 plants measured for five macronutrients and six micronutrients across a broad geographic gradient, provides a rare perspective on intraspecific trait variability in seed nutrient composition. To our knowledge, no comparable dataset exists that captures intraspecific seed nutrient variation at this spatial and ecological scale. Such breadth allows us to address patterns that remain largely unexplored, as previous studies have rarely examined seed nutrient composition across extensive geographic ranges within a wild species (but see De Frenne, Kolb, et al., 2011; Wang et al., 2025; Wu et al., 2024 for smaller-scale studies).

Seed nutrients were generally positively intercorrelated. The strongest correlation was between seed P and Mg (r=0.65, Figure 2), likely because their uptake and transport are partly coordinated (Weih et al., 2021), and because most seed P is stored as phytic acid, which binds cations including Mg (Wu et al., 2009). Overall, nutrient loading into seeds is often co-regulated during seed development (Himelblau & Amasino, 2001).

We found negative correlations between some seed nutrients, particularly Ca, K, and Ni, and the plant morphological traits, particularly seed weight. In other words, larger seeds had lower concentrations of these elements. Similar trade-offs between seed weight and protein content have been reported in cereals (Uauy et al., 2006) and the model species *Arabidopsis thaliana* (Chardon et al., 2014). This relationship has been linked to the accumulation of carbohydrates in the endosperm, specifically, larger seeds have more carbohydrates and thus lower concentrations of elemental nutrients (Sehgal et al., 2018).

The content of most elemental nutrients in seeds increased with mean annual temperature. This relationship was partially driven by decreasing seed weight towards warmer regions, particularly for seed elemental nutrients that negatively correlated with seed weight (Ca, K, and Ni). Plants from warmer areas had higher concentrations of these nutrients, likely because they had lower relative concentrations of carbohydrates. However, concentrations of nutrients that were unrelated to seed weight also increased with mean annual temperature (Mg, P, S, Mn, Ni, and Zn). Positive effect of mean annual temperature on nutrient content aligns with findings of a recent study on two grass species along a latitudinal gradient in China (Wang et al., 2025), and with a study on European forest herb (De Frenne, Kolb, et al., 2011). Nutrient uptake and transport in plant tissues are affected by temperature: as temperature increases towards optimum, uptake and transport of nutrients get faster (Hood & Mills, 1994; Mishra et al., 2023). This temperature-dependent mechanism could thus stand behind the increased contents of elemental nutrients in the seeds in warmer regions.

Soil characteristics had only a limited effect on nutrient concentration in the seeds. The strongest effect had soil pH, which negatively affected the concentration of Fe and Mn and positively affected Mo. The negative effects of pH on Fe and Mn are not surprising because the bioavailability of Fe and Mn is higher in acidic environments (Ramzani et al., 2016; Sims, 1986). The positive effect of pH on Mo is not as straightforward to explain because the uptake regulation of this element is complex (Kaiser et al., 2005). Surprisingly, soil available nutrients, specifically nitrate, phosphate, and sulphate, had no effect on the concentration of the respective elements in the seeds, and had, in general, only a very small effect on seed concentration of other elemental nutrients. This contrasts with ample literature on crops that shows that soil nutrient availability is crucial for nutrient concentrations in the seeds (Joy et al., 2015; Marschner, 2012; Wortmann et al., 2018). One possible explanation for the limited effect of soil properties on nutrient concentration in seeds can be the pooling of soil per population, which disabled us to capture fine-scale mosaics in soil characteristics experienced by individual plants. However, our results are in line with findings of the few existing studies on wild species that show that seed nutrient content across large geographic gradients is highly variable and only weakly correlates with soil properties (Wang et al., 2025; Wu et al., 2024).

### Population differentiation

While many of the environmental predictors had a significant effect on plant morphological traits and seed nutrient contents, they explained only 4.9-12.4 % and 3.3-22% of the variation, respectively. Much more variation (37-38 % and 24-57%, respectively) was explained by population identity, independent of environmental factors. In other words, plants growing in one population had more similar traits than predicted by environmental factors. One possible explanation is that our environmental predictors did not fully capture site-specific environments. However, the most important environmental drivers of plant traits across large geographic scales are associated with climate (Lemke et al., 2015), and thus, are unlikely to stand behind the large proportion of variability explained solely by the site identity. More likely, the similarity of plants growing at one site is caused by genetic factors. *H. murinum* is predominantly self-pollinating, and thus, most plants at one site are likely to be close relatives with identical or nearly identical genetic backgrounds (Volis et al., 2010). Possibly, also the variability associated with environmental predictors, particularly climate, might be at least partially genetically underpinned, because climatic gradients might covary with genetic differentiation(Leiblein-Wild & Tackenberg, 2014). However, as the present data were measured in-situ, it is impossible to disentangle the contribution of genetic variation and direct response to environment (plasticity) to variation in plant traits. This will require further research, including growing plants in a common environment, optimally in combination with molecular genetic analysis.

## Acknowledgements

We thank Johannes Kasper, Lea Klepka, Lena Lerbs, Laura Libera, Christina Mengel, Ann-Sophie Schmitt, Annika Sommerfeld, Sabine Ambrosius, and Maike Grusch for technical support and the CEPLAS Plant Metabolism and Metabolomics Laboratory Cologne for ICP-MS measurements.

