## Appendix for "Climate-driven in-situ trait variation in an annual ruderal grass across Europe"

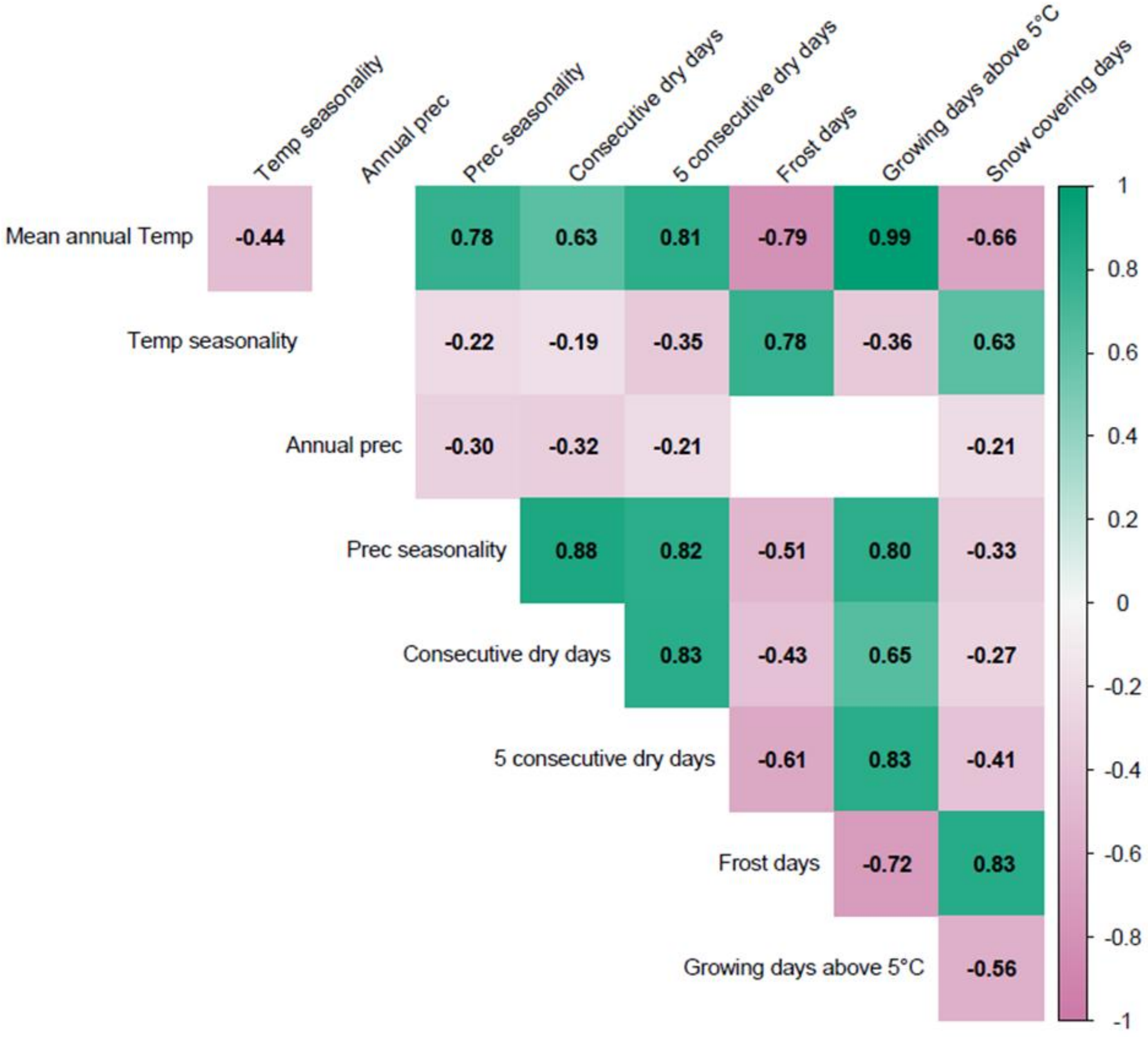


Figure S1| Pearson's correlation coefficient between all downloaded Chelsa climate variables available. Variables were averaged across all plants within each population. The number and color intensity in the cells indicate the strength and direction of two related variables. Empty cells indicate non-significant correlations (p > 0.05).

Table S1| Population differentiation in plant morphological traits and seed nutrient contents (macro- and micronutrients). Analysis of variance with error type I. Significant effects ($p < 0.05$) are highlighted in bold. $R^2$ values are variation explained by population differences, after accounting for the large number of sites in the analysis (adjusted $R^2$).

| **Traits** | Sum Sq | df | F | *p* | $R^2$ |
|---|---|---|---|---|---|
| *Morphology* | | | | | |
| Plant height | 907.50 | 204 | 7.21 | **< 0.001** | 0.38 |
| Seed weight | 863.13 | 199 | 6.96 | **< 0.001** | 0.38 |
| Spike number | 879.47 | 204 | 6.83 | **< 0.001** | 0.37 |
| *Seed macronutrients* | | | | | |
| Calcium (Ca) | 467.22 | 176 | 4.54 | **< 0.001** | 0.42 |
| Potassium (K) | 498.47 | 176 | 5.24 | **< 0.001** | 0.46 |
| Magnesium (Mg) | 551.34 | 176 | 6.73 | **< 0.001** | 0.53 |
| Phosphorus (P) | 461.39 | 176 | 4.42 | **< 0.001** | 0.41 |
| Sulfur (S) | 510.03 | 176 | 5.53 | **< 0.001** | 0.48 |
| *Seed micronutrients* | | | | | |
| Copper (Cu) | 576.80 | 176 | 7.64 | **< 0.001** | 0.57 |
| Iron (Fe) | 371.30 | 176 | 2.92 | **< 0.001** | 0.28 |
| Manganese (Mn) | 482.42 | 176 | 4.86 | **< 0.001** | 0.44 |
| Molybdenum (Mo) | 518.49 | 176 | 5.95 | **< 0.001** | 0.51 |
| Nickel (Ni) | 343.22 | 176 | 2.56 | **< 0.001** | 0.24 |
| Zinc (Zn) | 429.43 | 176 | 3.82 | **< 0.001** | 0.36 |

Table S2| Intraspecific variation in plant phenology and morphological traits, results of spatial lag model and linear mixed model (LMM), respectively, testing for the effect of climate and environmental factors, and bioavailable soil nutrients. Response variables were scaled. Analysis of variance for LMM with error type III. Significant effects (<0.05) are highlighted in bold.

| Plant traits | Predictor | Spatial units | SE | z | *p* | $R^2$ | |
|---|---|---|---|---|---|---|---|
| ***Phenology*** | *Spatial lag model* | | | | | | |
| Seed ripening | | 157 | | | | 0.71 | |
| (population mean) | Mean annual Temp | | 0.028 | -6.63 | **<0.001** | | |
| | Temp seasonality | | 0.045 | 2.00 | **0.045** | | |
| | Annual Prec | | 0.000 | 0.52 | 0.582 | | |
| | Soil pH | | 0.098 | -1.04 | 0.298 | | |
| | Vegetation cover | | 0.003 | 0.52 | 0.603 | | |
| | Soil nitrate | | 0.012 | 3.16 | **0.002** | | |
| | Soil phosphate | | 0.131 | 0.08 | 0.939 | | |
| | Soil sulfate | | 0.136 | -1.28 | 0.200 | | |
| | | df | Resid df | F | *p* | $R^2$ (fixed) | $R^2$ (random) |
| ***Morphology*** | *Linear mixed model* | | | | | | |
| Plant height | | | | | | 0.12 | 0.33 |
| (log) | Mean annual Temp | 1 | 166.4 | 5.33 | **0.022** | | |
| | Temp seasonality | 1 | 165.3 | 0.27 | 0.607 | | |
| | Annual Prec | 1 | 165.2 | 10.92 | **0.001** | | |
| | Soil pH | 1 | 164.1 | 1.88 | 0.172 | | |
| | Vegetation cover | 1 | 1705.7 | 164.53 | **<0.001** | | |
| | Soil nitrate | 1 | 168.4 | 0.00 | 0.998 | | |
| | Soil phosphate | 1 | 181.2 | 0.01 | 0.932 | | |
| | Soil sulfate | 1 | 193.6 | 0.21 | 0.644 | | |
| Seed weight | | | | | | 0.09 | 0.32 |
| | Mean annual Temp | 1 | 166.1 | 4.14 | **0.044** | | |
| | Temp seasonality | 1 | 164.8 | 0.00 | 0.984 | | |
| | Annual Prec | 1 | 164.6 | 6.40 | **0.012** | | |
| | Soil pH | 1 | 164.1 | 0.20 | 0.655 | | |
| | Vegetation cover | 1 | 1679.0 | 89.66 | **<0.001** | | |
| | Soil nitrate | 1 | 167.8 | 0.01 | 0.904 | | |
| | Soil phosphate | 1 | 179.8 | 0.14 | 0.708 | | |
| | Soil sulfate | 1 | 192.3 | 2.07 | 0.152 | | |
| Spike number | | | | | | 0.05 | 0.34 |
| (log) | Mean annual Temp | 1 | 166.2 | 8.35 | **0.004** | | |
| | Temp seasonality | 1 | 165.2 | 0.33 | 0.568 | | |
| | Annual Prec | 1 | 165.1 | 0.64 | 0.426 | | |
| | Soil pH | 1 | 164.1 | 0.22 | 0.642 | | |
| | Vegetation cover | 1 | 1710.6 | 6.52 | **0.011** | | |
| | Soil nitrate | 1 | 167.7 | 0.62 | 0.431 | | |
| | Soil phosphate | 1 | 180.0 | 0.09 | 0.768 | | |
| | Soil sulfate | 1 | 191.6 | 0.06 | 0.809 | | |

Table S3| Intraspecific variation in seed macronutrient content, results of linear mixed model testing for the effect of climate and environmental factors, and bioavailable soil nutrients. Response variables were scaled. Analysis of variance with error type III. Significant effects (<0.05) are highlighted in bold.

| Seed macronutrients | Predictor | df | Resid df | F | *p* | $R^2$ (fixed) | $R^2$ (random) |
|---|---|---|---|---|---|---|---|
| Calcium (Ca) | | | | | | 0.10 | 0.32 |
| (log) | Mean annual Temp | 1 | 144.9 | 13.53 | **<0.001** | | |
| | Temp seasonality | 1 | 144.1 | 1.30 | 0.256 | | |
| | Annual Prec | 1 | 142.4 | 1.70 | 0.194 | | |
| | Soil pH | 1 | 150.0 | 3.52 | 0.062 | | |
| | Vegetation cover | 1 | 782.4 | 8.10 | **0.005** | | |
| | Soil nitrate | 1 | 145.7 | 0.07 | 0.798 | | |
| | Soil phosphate | 1 | 145.0 | 0.00 | 0.975 | | |
| | Soil sulfate | 1 | 150.5 | 0.10 | 0.750 | | |
| Potassium (K) | | | | | | 0.12 | 0.37 |
| (log) | Mean annual Temp | 1 | 145.6 | 7.95 | **0.005** | | |
| | Temp seasonality | 1 | 144.9 | 6.44 | **0.012** | | |
| | Annual Prec | 1 | 143.4 | 4.51 | **0.035** | | |
| | Soil pH | 1 | 150.0 | 3.39 | 0.068 | | |
| | Vegetation cover | 1 | 781.4 | 7.75 | **0.006** | | |
| | Soil nitrate | 1 | 146.6 | 0.45 | 0.504 | | |
| | Soil phosphate | 1 | 145.8 | 0.05 | 0.831 | | |
| | Soil sulfate | 1 | 151.7 | 0.94 | 0.334 | | |
| Magnesium (Mg) | | | | | | 0.22 | 0.34 |
| | Mean annual Temp | 1 | 145.8 | 65.76 | **<0.001** | | |
| | Temp seasonality | 1 | 145.1 | 12.48 | **0.001** | | |
| | Annual Prec | 1 | 143.6 | 4.46 | **0.036** | | |
| | Soil pH | 1 | 150.0 | 2.24 | 0.137 | | |
| | Vegetation cover | 1 | 780.0 | 1.46 | 0.227 | | |
| | Soil nitrate | 1 | 146.8 | 4.34 | **0.039** | | |
| | Soil phosphate | 1 | 146.0 | 0.60 | 0.441 | | |
| | Soil sulfate | 1 | 152.1 | 0.01 | 0.904 | | |
| Phosphorus (P) | | | | | | 0.10 | 0.30 |
| (log) | Mean annual Temp | 1 | 144.6 | 20.75 | **<0.001** | | |
| | Temp seasonality | 1 | 143.8 | 2.55 | 0.112 | | |
| | Annual Prec | 1 | 142.1 | 0.33 | 0.566 | | |
| | Soil pH | 1 | 150.0 | 3.59 | **0.060** | | |
| | Vegetation cover | 1 | 780.4 | 0.68 | 0.411 | | |
| | Soil nitrate | 1 | 145.3 | 6.86 | **0.010** | | |
| | Soil phosphate | 1 | 144.6 | 2.15 | 0.145 | | |
| | Soil sulfate | 1 | 150.0 | 0.54 | 0.465 | | |
| Sulfur (S) | | | | | | 0.08 | 0.39 |
| | Mean annual Temp | 1 | 145.6 | 15.64 | **<0.001** | | |
| | Temp seasonality | 1 | 145.0 | 0.03 | 0.867 | | |
| | Annual Prec | 1 | 143.5 | 1.28 | 0.260 | | |
| | Soil pH | 1 | 150.0 | 2.45 | 0.119 | | |
| | Vegetation cover | 1 | 781.0 | 0.64 | 0.426 | | |
| | Soil nitrate | 1 | 146.7 | 2.19 | 0.141 | | |
| | Soil phosphate | 1 | 145.9 | 0.06 | 0.810 | | |
| | Soil sulfate | 1 | 151.9 | 1.55 | 0.215 | | |

Table S4| Intraspecific variation in seed micronutrient content, results of linear mixed model testing for the effect of climate and environmental factors, and bioavailable soil nutrients. Response variables were scaled. Analysis of variance with error type III. Significant effects (< 0.05) are highlighted in bold.

| Seed micronutrient | Predictor | df | Resid df | F | *p* | $R^2$ (fixed) | $R^2$ (random) |
|---|---|---|---|---|---|---|---|
| Copper (Cu) | | | | | | 0.12 | 0.45 |
| | Mean annual Temp | 1 | 146.8 | 0.41 | 0.523 | | |
| | Temp seasonality | 1 | 146.1 | 9.10 | **0.003** | | |
| | Annual Prec | 1 | 145.0 | 0.01 | 0.935 | | |
| | Soil pH | 1 | 150.0 | 0.43 | 0.513 | | |
| | Vegetation cover | 1 | 764.5 | 5.49 | **0.019** | | |
| | Soil nitrate | 1 | 148.2 | 1.17 | 0.281 | | |
| | Soil phosphate | 1 | 147.1 | 0.05 | 0.832 | | |
| | Soil sulfate | 1 | 154.1 | 6.35 | **0.013** | | |
| Iron (Fe) | | | | | | 0.03 | 0.27 |
| (log) | Mean annual Temp | 1 | 143.9 | 0.00 | 0.969 | | |
| | Temp seasonality | 1 | 142.8 | 1.52 | 0.219 | | |
| | Annual Prec | 1 | 141.0 | 0.15 | 0.696 | | |
| | Soil pH | 1 | 149.8 | 5.18 | **0.024** | | |
| | Vegetation cover | 1 | 766.8 | 1.64 | 0.200 | | |
| | Soil nitrate | 1 | 144.2 | 1.97 | 0.162 | | |
| | Soil phosphate | 1 | 143.4 | 0.00 | 0.948 | | |
| | Soil sulfate | 1 | 148.6 | 0.00 | 0.960 | | |
| Manganese (Mn) | | | | | | 0.10 | 0.34 |
| | Mean annual Temp | 1 | 145.1 | 8.84 | **0.003** | | |
| | Temp seasonality | 1 | 144.4 | 2.81 | 0.096 | | |
| | Annual Prec | 1 | 142.8 | 1.16 | 0.284 | | |
| | Soil pH | 1 | 150.0 | 24.22 | **<0.001** | | |
| | Vegetation cover | 1 | 783.0 | 0.01 | 0.903 | | |
| | Soil nitrate | 1 | 146.0 | 2.66 | 0.105 | | |
| | Soil phosphate | 1 | 145.3 | 0.03 | 0.861 | | |
| | Soil sulfate | 1 | 150.9 | 0.78 | 0.378 | | |
| Molybdenum (Mo) | | | | | | 0.08 | 0.42 |
| (sqrt) | Mean annual Temp | 1 | 148.3 | 7.10 | **0.009** | | |
| | Temp seasonality | 1 | 146.6 | 0.51 | 0.477 | | |
| | Annual Prec | 1 | 147.0 | 3.91 | **0.050** | | |
| | Soil pH | 1 | 160.2 | 6.13 | **0.014** | | |
| | Vegetation cover | 1 | 753.4 | 0.29 | 0.593 | | |
| | Soil nitrate | 1 | 146.9 | 2.14 | 0.146 | | |
| | Soil phosphate | 1 | 146.9 | 0.41 | 0.522 | | |
| | Soil sulfate | 1 | 151.9 | 2.94 | 0.088 | | |
| Nickel (Ni) | | | | | | 0.09 | 0.20 |
| (log) | Mean annual Temp | 1 | 143.2 | 22.14 | **<0.001** | | |
| | Temp seasonality | 1 | 141.6 | 12.72 | **<0.001** | | |
| | Annual Prec | 1 | 139.8 | 5.23 | **0.024** | | |
| | Soil pH | 1 | 149.5 | 1.48 | 0.225 | | |
| | Vegetation cover | 1 | 738.3 | 17.43 | **<0.001** | | |
| | Soil nitrate | 1 | 142.7 | 3.98 | **0.048** | | |
| | Soil phosphate | 1 | 141.9 | 0.07 | 0.785 | | |
| | Soil sulfate | 1 | 147.0 | 0.89 | 0.347 | | |
| Zinc (Zn) | | | | | | 0.05 | 0.31 |
| (log) | Mean annual Temp | 1 | 144.5 | 4.87 | **0.029** | | |
| | Temp seasonality | 1 | 143.6 | 0.00 | 0.953 | | |
| | Annual Prec | 1 | 141.8 | 0.08 | 0.776 | | |
| | Soil pH | 1 | 149.8 | 2.97 | 0.087 | | |
| | Vegetation cover | 1 | 776.1 | 4.17 | **0.042** | | |
| | Soil nitrate | 1 | 145.0 | 1.46 | 0.230 | | |
| | Soil phosphate | 1 | 144.4 | 2.05 | 0.154 | | |

| | | | | |
|---|---|---|---|---|
| Soil sulfate | 1 | 149.6 | 0.90 | 0.343 |

### *Appendix*

*Ploidy levels in Hordeum murinum – methods*

For the ploidy test, we included the majority of populations, except 16 populations from the northern part of the distribution that were less than 10 km apart from another population, because we expected and later confirmed that there are only tetraploids in this region. Given the large number of samples to be tested (in total, more than 2300 plants), we implemented a stratified approach. In the first step, we screened all populations to detect regions where both ploidies occur. To do this, we analyzed one random plant per population. We detected the presence of diploids only in southern Europe. Thus, for all populations south of 47°, we then analyzed six to ten random individual plants pooled into one sample. If this procedure indicated the presence of both ploidy levels within one population, we verified this by analyzing each of the six to ten plants individually.

We measured the ploidy using a flow cytometer (CyFlow™ Ploidy Analyser, Sysmex Europe SE). To do this, we sowed the seeds from sampled mother plants in quick pots, grew seedlings, and collected approximately 1 cm leaf samples. We placed the leaf samples in a 5.5 cm diameter petri dish and chopped them with a razor into small pieces with two drops of CyStain™ UV Ploidy staining solution (Sysmex Europe SE, Norderstedt, Germany). After chopping, we added 2 mL of staining solution to the leaf mass and transferred the mixture to 3.5 mL sample tubes fitted with a CellTrics 50 µm filter mesh (Sysmex Europe SE, Norderstedt, Germany) to remove plant debris, leaving only the filtered cell suspension. Before we loaded the sample to the flow cytometer, we loaded reference samples (diploid, *H. murinum glaucum* acc. BCC2017, which was regenerated from NGB90348, and tetraploid, *H. murinum murinum* acc. BCC2009) to determine the peak position of both ploidy levels. With the pre-installed Windows™-based FCM software CyView 1.8 (Sysmex Europe SE, Norderstedt, Germany), we were able to perform real-time data display and analysis. Each sample was classified as either diploid or tetraploid based on the similarity of the projected peaks to the reference samples with confirmed ploidy levels.

*Geographic distribution of ploidy levels*

We found that diploid plants (*H. murinum* subsp. *glaucum*) were exclusively present in Spain (mainland), Morocco, and Algeria in our collection (Figure S2). From the 47 populations sampled in this region, 16 were only tetraploids, 11 were only diploids, and 20 populations had mixed ploidy (Figure S2). For the main study, we retained only the 16 fully tetraploid populations from this region.

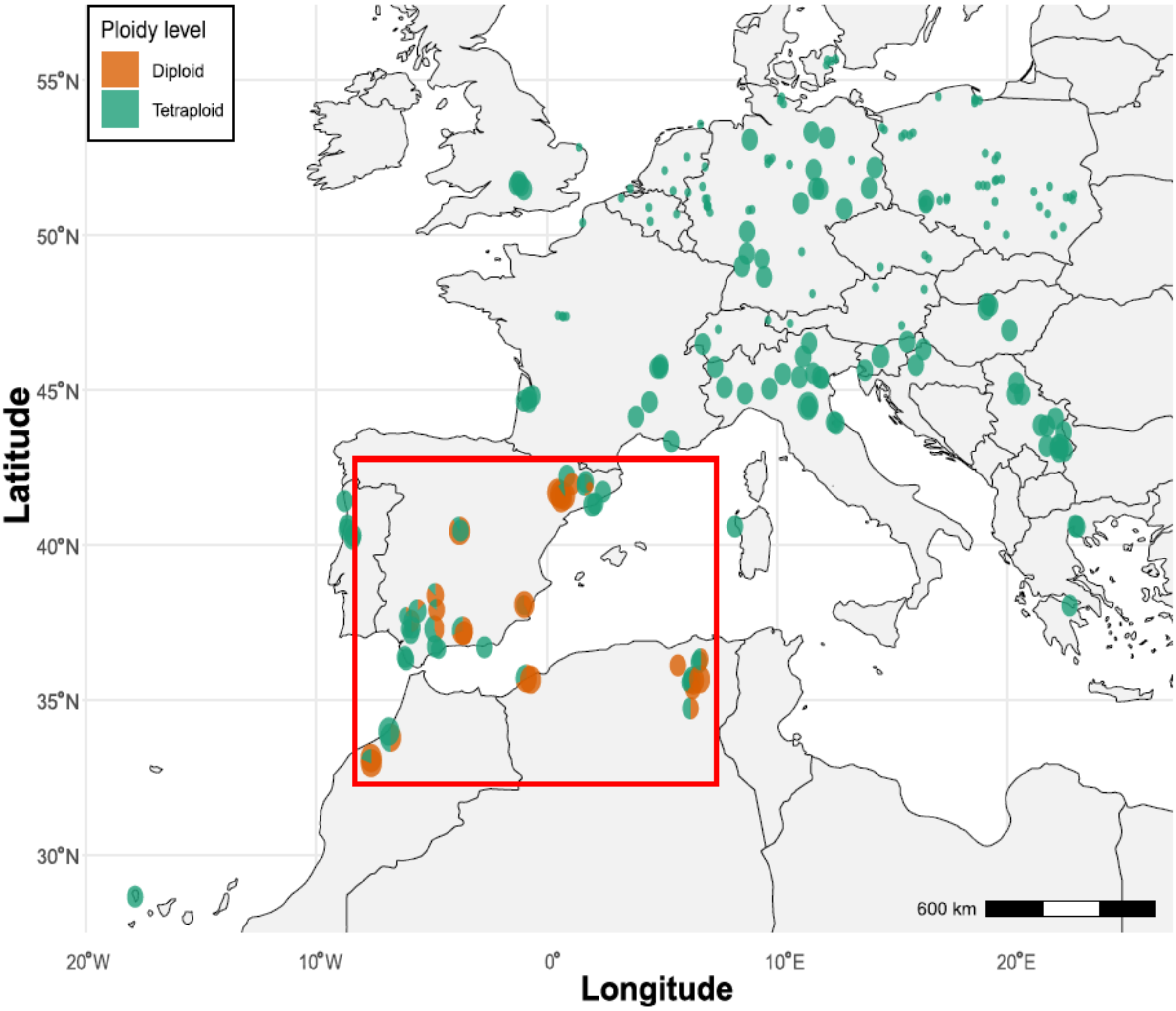


Figure S2| Ploidy level of tested individuals of sampled *Hordeum murinum* populations across the European range. Point sizes indicate the number of tested mother plants per population (1-10 tested individuals), and pie charts exhibit mixed ploidy levels within a population. The red square displays the regions with both recorded ploidy levels. Diploids are colored in orange and tetraploids in green.

*Trait differences between ploidy levels*

To test for in-situ differences between diploids and tetraploids, we focused only on populations from the region where both ploidies grew next to each other (red square in Figure S2). We related plant morphological traits (plant height, spike number, and seed weight) and seed nutrient contents (macro- and micronutrients) as response variables to ploidy level as the fixed effect and population identity as the random effect factor, in a linear mixed model effect using the *lme4* package (Bates et al., 2015). We found that tetraploids had heavier seeds. We found no differences between ploidy levels in any other trait (Table S5).

Table S5| Differences between the two ploidy levels in plant morphological traits and seed nutrient content (macro- and micronutrients). Response variables were scaled, results from a linear mixed model. Analysis of variance with error type II. Significant effects ($p < 0.05$) are highlighted in bold.

| | *Ploidy level* | | | | | |
|---|---|---|---|---|---|---|
| **Trait** | Estimate | Std. error | Df | Resid. Df | *F* | *p* |
| *Plant morphological traits* | | | | | | |
| Plant height | -0.0004 | 0.12 | 1 | 323.9 | 0.00001 | 0.997 |
| Seed weight | 0.91 | 0.13 | 1 | 245.4 | 49.20 | **<0.001** |
| Spike number | -0.19 | 0.10 | 1 | 332.2 | 3.28 | 0.071 |
| *Seed macronutrients* | | | | | | |
| Calcium (Ca) | -0.26 | 0.23 | 1 | 114.1 | 1.26 | 0.264 |
| Magnesium (Mg) | 0.01 | 0.24 | 1 | 67.7 | 0.0001 | 0.979 |
| Potassium (K) | 0.31 | 0.20 | | 120.0 | 2.38 | 0.126 |
| Phosphorus (P) | 0.14 | 0.24 | 1 | 86.5 | 0.35 | 0.554 |
| Sulfur (S) | -0.08 | 0.23 | 1 | 115.9 | 0.12 | 0.729 |
| Seed micronutrients | | | | | | |
| Copper (Cu) | -0.14 | 0.24 | 1 | 106.8 | 0.31 | 0.579 |
| Iron (Fe) | -0.30 | 0.24 | 1 | 109.5 | 1.48 | 0.227 |
| Manganese (Mn) | -0.39 | 0.22 | 1 | 117.4 | 3.00 | 0.086 |
| Molybdenum (Mo) | -0.44 | 0.25 | 1 | 86.4 | 3.03 | 0.086 |
| Nickel (Ni) | -0.22 | 0.23 | 1 | 53.4 | 0.87 | 0.356 |
| Zinc (Zn) | -0.30 | 0.24 | 1 | 100.3 | 1.52 | 0.221 |

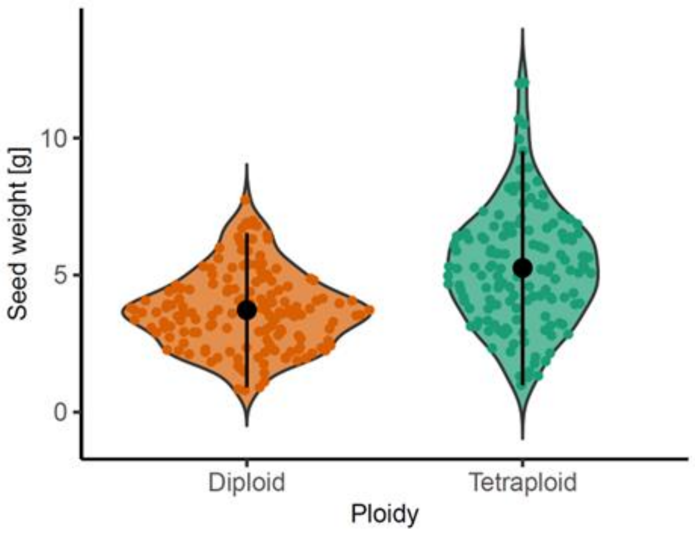


Figure S3| Differences between diploids and tetraploids in seed weight ($p<0.001$). Each point represents an individual, with colors indicating ploidy levels: diploid in orange, and tetraploid in green.

*Habitat preferences of ploidy levels*

To illustrate niche differences between the two ploidy levels (diploid (2x) and tetraploid (4x)), we performed a PCA (principal component analysis) with the climatic and environmental factors (Figure S4), and colored the individual plants according to their ploidy level.

The first two principal component axes, PC1 and PC2, explained 30.61% and 18.95% of the total variance, respectively. PC1 was strongly positively associated with annual precipitation and negatively with temperature seasonality, soil sulfate, and soil pH. PC2 was associated with soil nitrate and mean annual temperature. We found partial clustering of diploids and tetraploids, particularly along the PC1 axis. Tetraploids were partially associated with higher annual precipitation, lower pH and potentially lower temperature seasonality. In contrast, diploids were predominantly associated with higher temperature seasonality, higher soil sulfate, phosphate, and pH levels, but lower annual precipitation (Figure S4).

This visualization indicates that diploids tend to occur in slightly drier environments and higher temperature seasonality, while tetraploids are more frequent in regions with higher precipitation.

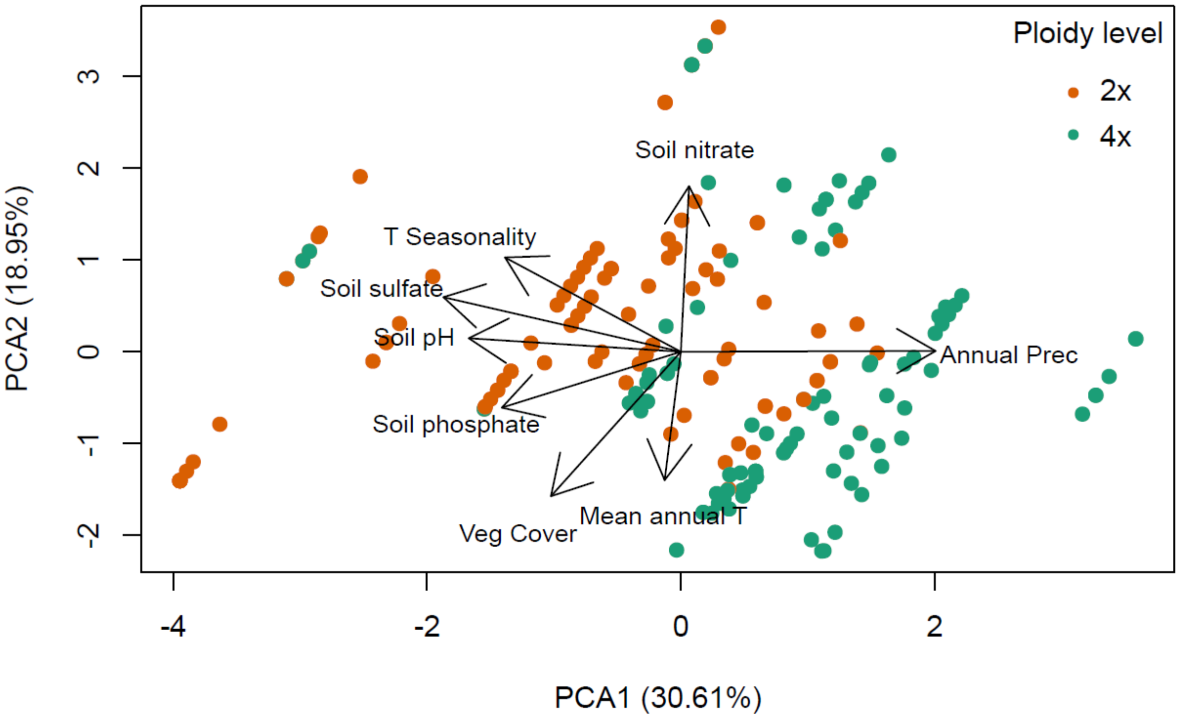


Figure S4| Ploidy level distribution linked with climatic and environmental conditions. Each point represents an individual, with colors indicating ploidy levels: diploid (2x) in orange, and tetraploid (4x) in green. Climatic and environmental variables (black arrows) indicate the direction and strength of their contribution to the principal components. Longer arrows denote stronger correlations with the respective axes.
